# Sedimentary DNA can reveal the past population dynamics of a pelagic copepod

**DOI:** 10.1101/2022.09.09.506697

**Authors:** Kai Nakane, Xin Liu, Hideyuki Doi, Gaël Dur, Michinobu Kuwae, Syuhei Ban, Narumi Tsugeki

## Abstract

1. Copepods play a key trophic role as secondary producers, transferring primary production to higher trophic levels such as fish. Copepod production contributes significantly to successful fish recruitment. Despite their importance, knowledge of copepod dynamics over several decades remains limited due to the lack of long-term data series with adequate sampling and analysis. However, an understanding of long-term copepod dynamics is urgently required to strive toward better management for sustainable aquatic ecosystems and fish recruitment.
2. Sedimentary DNA (sedDNA) has been developing as a useful tool for reconstructing past plankton dynamics. This study evaluates whether sedDNA targeting the pelagic copepod, *Eodiaptomus japonicus*, in Lake Biwa (Japan) can be an effective tool for elucidating its past population dynamics.
3. We applied a quantitative polymerase chain reaction method targeting the mitochondrial cytochrome c oxidase subunit I gene on two sediment cores and compared the detected sedDNA concentrations with the unique long-term dataset of demographic traits, biomass, specific growth rate, production, subitaneous eggs, and resting eggs of *E. japonicus*.
4. The sedDNA concentration of *E. japonicus* recovered from sediment layers correlated significantly with *in situ* production, biomass, and production of immediately hatching eggs but not with resting eggs or specific growth rate. Our study provides evidence for the effective use of sedDNA as a tracking tool for assessing past copepod production dynamics.

## Introduction

Copepods are the most abundant multicellular organisms on Earth (Schminke, 2007) and are important secondary producers that transfer primary production to higher trophic levels such as to fish in aquatic ecosystems (Mauchline, 1998). Copepod production as a food resource is essential for sustainable fish management (Mauchline, 1998). Successful fish recruitment is highly dependent on copepod production because it promotes fish larval survival (Mauchline, 1998). Copepod population growth is influenced by temperature and food availability (Ban, 1994; Hart, 1990; Koski & Kuosa, 1999; Liu *et al*., 2015). Previous studies have shown that global warming decreases copepod abundance (Garzke *et al*., 2015). Also, large-scale climate events showing oscillation cycles over decades, such as the Pacific Decadal Oscillation and Arctic Oscillation, regulate ocean temperatures and consequently affect copepod biomass and production (Chiba *et al*., 2006; Liu *et al*., 2021; Peterson & Schwing, 2003). Information about long-term copepod population dynamics is urgently required to achieve better fish management and to understand how large-scale climatic events influence aquatic ecosystems. Nevertheless, copepod dynamics across decades are rarely investigated due to the cost and time required to obtain long-term data series.

Sedimentary archives are valuable for gaining information on zooplankton long-term dynamics (Jankowski & Straile, 2003; Brede *et al*., 2009). For specific species, such as the cladoceran zooplankton *Daphnia*, whose remains and resting eggs are well preserved in sediment, morphological investigation and genetic analysis are powerful tools to clarify biological responses to anthropogenic perturbations (Brede *et al*., 2009; Jankowski & Straile, 2003; Monchamp *et al*., 2017; Tsugeki *et al*., 2009). However, copepods have been excluded from palaeoecological studies because their remains are not well preserved in sediments (Selden *et al*., 2010). Although copepod resting eggs are preserved in sediments (Hairston *et al*., 1995), they are seldom used in palaeoecological studies because of their poor taxonomic features (Briski *et al*., 2011). Consequently, traditional investigations of sediment archives are insufficient for establishing past copepod dynamics.

To overcome such difficulties, sedimentary DNA (sedDNA) can be a useful proxy for reconstructing past dynamic of organisms without visible remains (Capo *et al*., 2021). Through the sedDNA approach, great progress has been made in reconstructing the past dynamics of various organisms such as mammals (Giguet-Covex *et al*., 2014), plants (Alsos *et al*., 2018; Epp *et al*., 2015; Giguet-Covex *et al*., 2014), plankton (Epp *et al*., 2010, 2015; Huang *et al*., 2020; Monchamp *et al*., 2016) and fish (Kuwae *et al*., 2020). Combined studies of sedDNA with morphological techniques have been conducted to track plankton community composition (Huang *et al*., 2020; Monchamp *et al*., 2016). However, most studies on plankton sedDNA have focused on phytoplankton; thus, the utility of the sedDNA approach to reconstruct zooplankton dynamics still remains limited. A recent study of zooplankton sedDNA targeting *Daphnia galeata* and *D. pulicaria* in Lake Biwa (Japan) using quantitative polymerase chain reaction (qPCR) successfully detected the sedDNA of both *Daphnia* species (Tsugeki *et al*., 2022). The temporal variation in *Daphnia* sedDNA concentration was consistent with their resting egg production, indicating that zooplankton sedDNA could be used as a tool for monitoring their spawning activity (Tsugeki *et al*., 2022). Previous studies also successfully sequenced copepod DNA in sediment samples (Bissett *et al*., 2005; Epp *et al*., 2015; Xu *et al*., 2011) and reconstructed past variation in the copepod community of a North Greenland lake (Epp *et al*., 2015) and in Antarctic lakes (Bissett et al. 2005). However, copepod DNA was only retrieved from a limited number of samples, perhaps due to the relatively long fragment amplified by the available primers (as previously pointed out Epp *et al*. 2015). These findings suggest that historically deposited copepod DNA is sufficiently well preserved in sediments to allow for reconstruction of long-term population and community level changes. Nevertheless, the ability to detect the sedDNA of a specific copepod species and to quantitatively reconstruct its dynamics is limited. Thus, there is an urgent need to develop specific sedDNA techniques for the quantitative reconstruction of the dynamics key individual copepod taxa.

To ensure the reliability of using sedDNA to reconstruct past population dynamics, accumulating knowledge about the source of sedDNA is imperative. Without such information, one cannot interpret what is being reconstructed. The comparison of historical variations between sedDNA and regular survey data in the water column could confirm the reliability of the information provided by sedDNA. In Lake Biwa (Japan), one of the world’s most ancient lakes (Hampton *et al*., 2018), regular zooplankton surveys have been conducted intensively since 1965 (Hsieh *et al*., 2011; Liu *et al*., 2020b). Moreover, the long-term demographic traits of the dominant pelagic copepod, *Eodiaptomus japonicus*, such as biomass, population growth rate, and production, were determined from 1971 to 2010 using experimental and microscopic observations (Liu *et al*., 2015, 2020b, 2021). *Eodiaptomus japonicus* produces both subitaneous eggs (that hatch within a few days, without requiring a period of dormancy) and resting eggs (which require a period of dormancy before hatching); the historical variation in resting egg concentration was recovered from sediment archives up to 60 years old (Liu *et al*., 2020a). Sediment cores from Lake Biwa can provide high-resolution temporal data and exhibit fairly constant sedimentation rates (2–4 years per cm) over this time frame (Tsugeki et al., 2003, 2021). Therefore, *E. japonicus* is a suitable candidate for comparing historical variation between its sedDNA and demographic traits over the past several decades.

We first investigated whether sedDNA of *E. japonicus* could be successively and quantitatively detected from sediment core samples collected in the deep and large northern basin of Lake Biwa via qPCR analysis. We then tested the efficiency of sedDNA as a tool to reconstruct quantitative population dynamics (Fig. 1b) by comparing the detected *E. japonicus* sedDNA with its previously determined demographic traits (Fig. 1g, h). To evaluate the contribution of the DNA derived from resting eggs to the sedDNA sample, we conducted a sieving experiment (Fig. 1e). In addition, we also investigated the sedDNA concentrations in pore water separated from the residual sediment to clarify the influence of its vertical movement (Fig. 1f). Furthermore, we assessed the effects of enzymatic inhibitors in the qPCR process (Fig. 1b), and the effects of the co-precipitation with phytoplankton debris and their aggregates (Fig. 1d). Based on the findings, we discuss what was reconstructed for *E. japonicus* from sedDNA and its reliability as a tool for reconstructing *in situ E. japonicus* population dynamics.

**Figure 1.**
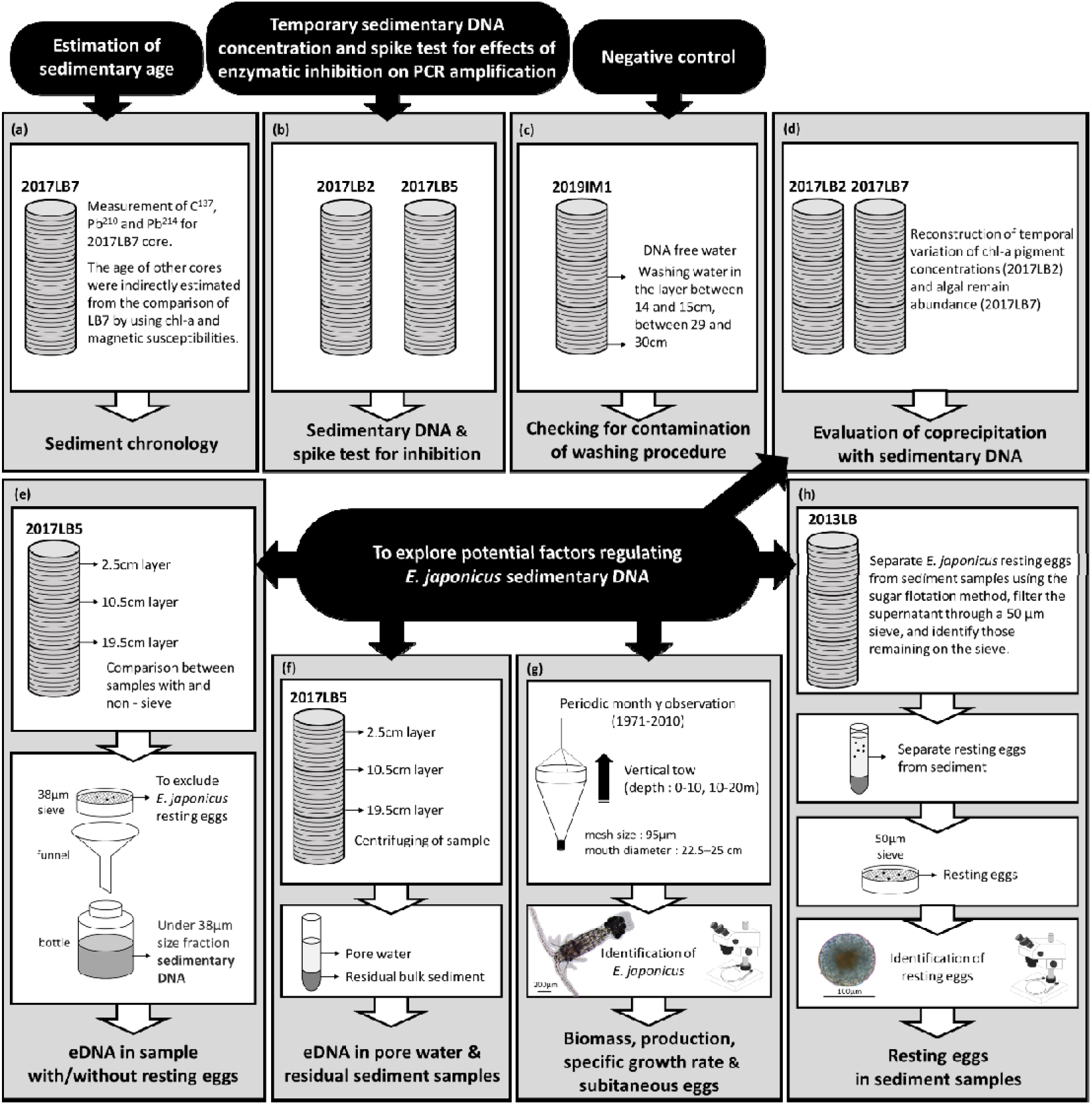
Research and experimental design of the study. (a) Sediment geochronology; (b) Reconstruction of temporal variation in *Eodiaptomus japonicus* sedimentary DNA (sedDNA) concentrations and spike test for inhibition; (c) Checking for contamination; (d) Reconstruction of temporal variation of chlorophyll-a pigment concentration and algal remains abundance; (e) Evaluating potential sources of *E. japonicus* sedDNA using the sieved and unsieved samples; (f) Regulating factors of *E. japonicus* sedDNA; (g) Demographic traits of biomass, specific growth rate, production, and subitaneous eggs of *E. japonicus*; (h) Sedimentary records of resting eggs. The named LB cores were collected in November 2013 (2013LB) and August 2017 (2017LB2, 2017LB5, and 2017LB7), and IM core (namely 2019IM1) was collected in August 2019 (details are described in the section on sampling site and sediment collection of methods).

## Methods

### Sampling site, sediment core, and zooplankton sample collection

We used sediment core samples to reconstruct historical variation in sedDNA concentrations of *E. japonicus* at Lake Biwa (Fig. 1b). We collected a total of four 30-cm long gravity core samples (Fig. 1a-f) from two points in the northern basin of Lake Biwa (Fig. S1). Three were collected on August 17, 2017 (2017LB core: 2017LB2, 2017LB5, 2017LB7; 35°15′1.0″N, 136°04′0.78″ E; water depth = 71 m), and one on August 27, 2019 (2019IM core: 2019IM1; 35°22′37″N, 136°05′48″E; water depth = 93 m). The sampling methods have been described in detail previously (Tsugeki *et al*., 2022). The 2017LB2 and 2017LB5 cores were used to determine the variation in *E. japonicus* sedDNA concentrations (Fig. 1b, Table S2) and to explore potential sources of its sedDNA (Fig. 1e, f). The 2017LB7 core was used for the chronology and for some algal remains whereas the 2019IM1 core was used as the negative control (details are described below).

The collected cores were then cut into successive sections each 1 cm thick using a vertical extruder with a cutting apparatus. After sectioning, each sliced sample was homogenized by hand shaking and then, all subsamples were taken from each homogenized sample. Between sample preparations, all pipes, knives, and cutting apparatuses were rinsed with 0.6% sodium hypochlorite, tap water, and Milli-Q water to prevent DNA cross-contamination. All sliced samples were transferred to lightproof bags and frozen at −80°C until further analysis. To assess the level of potential contamination during the core cutting, DNA extraction, and qPCR analysis processes, we inserted control water samples at a depth of 14-15 cm and 29-30 cm layers; hereafter expressed as the middle depth, i.e., 14.5 cm and 29.5 cm, respectively, in the sediment cores. We used the water samples for core 2019IM1 as the negative control (Fig. 1c).

To develop a primer-probe specific for *E. japonicus*, we collected a total of 95 individuals (including 14 egg-bearing individuals) and 91 individuals of the sympatric copepod *Mesocyclops* spp. (including 17 egg-bearing individuals) from water samples on May 14, 2021, in Lake Biwa. Among these, six individuals of *E. japonicus* (including three egg-bearing individuals) and four individuals of *Mesocyclops* spp. (including two egg-bearing individuals) were randomly selected, stored at –80 L until DNA extraction, and then stored at –20 L until *in vitro* validation.

### Chronology of sediment cores

Details of the chronological method have been reported elsewhere (Tsugeki *et al*., 2021). Briefly, sediment chronology was established for the 2017LB7 core based on the constant rate of supply (CRS) model of ^210^Pb dating (Appleby & Oldfield, 1978). Results were verified using the ^137^Cs peak traced in the period 1962 to 1963 (Appleby, 2001). The ages and age errors of the 2017LB2 and 2017LB5 cores were estimated indirectly, making use of stratigraphic correlations between the cores based on chronological controls in chlorophyll pigments and magnetic susceptibilities of the chronological 2017LB7 core (Tsugeki *et al*., 2021). The marked peak or trough layers were used as reference layers to compare these proxies (Fig. S3). In this study, sedimentary records of resting eggs for *E. japonicus* (Liu *et al*., 2020a) were newly denoted with improved chronology estimated using the CRS model (Fig. S5a, b).

### DNA extraction and primer-probe development for *E. japonicus*

DNA extraction and purification of a total of 61 sediment samples for two cores (Table S2; 2017LB2: 34 samples, 2017LB5: 27 samples) followed the methodology described in previous papers (Kuwae *et al*., 2020; Tsugeki *et al*., 2022; Sakata *et al*., 2020). This involved using the ‘Experienced User Protocol 3 to 22’ of a fecal-soil DNA extraction kit (Power Soil DNA Isolation Kit, Qiagen, Germany). In short, 9 g of each sample was incubated in a 9 ml alkaline solution (comprising 6 ml of 0.33 M sodium hydroxide and a 3 mL Tris-EDTA buffer (pH 6.7)) in a 45 ml tube at 94 °C for 50 min. After incubation, samples were centrifugation at 10,000×g at 4 °C for 60 min. We took 7.5 mL of the supernatant of the alkalized mixture and transferred it to a new tube and neutralized the solution with 7.5 mL of 1 M Tris–HCl (pH 6.7). After adding 1.5 mL of 3 M sodium acetate (pH 5.2) and 30 mL ethanol (99.8%), the solution was stored at −20 °C for more than 1 h and then centrifuged at 10,000×g at 4 °C for 60 min. The supernatant was decanted. The residual pellet was mixed with 100 μL of DNA-free water and transferred to a ‘PowerBead Tube’ that was installed in a PowerSoil DNA kit. Finally, elution was in 200 µL of the DNA solution and stored at –20 L until qPCR analysis.

The primer-probe for *E. japonicus* in qPCR analysis was designed for this study as follows. Firstly, we downloaded the mitochondrial cytochrome c oxidase subunit I gene (COI) from the National Center for Biotechnology Information (NCBI, http://www.ncbi.nlm.nih.gov/). We then compared the COI genes of the target species (*E. japonicus*) with those of the non-target copepod species *Mesocyclops thermocyclopoides*, *Cyclops vicinus, Daphnia galeata*, and *D. pulicaria* (Table S1b) which are the main zooplankton species in Lake Biwa (Liu *et al*., 2020b). The reference sequences for the targeted gene regions were queried on possible amplicons between 50 and 150 bp using NCBI primer-BLAST (http://www.ncbi.nlm.nih.gov/tools/primer-blast/), after which we constructed the primer-probe using Primer3plus (https://www.bioinformatics.nl/cgi-bin/primer3plus/primer3plus.cgi). The primer-probe was designed as follows (Table S1a): forward primer, 5′-GTC CCT TAA TCG ATA GTC CTC G-3′; reverse primer, 5′-TCA CCT CCA CCC CCA TAT AA-3′; and probe sequence, 5′-FAM-AAC TTC AGG TCA AGG TGC AG-TAMRA-3′. The specificities of the primers and probe were then confirmed *in silico* with homologous sequences from other copepods and *Daphnia* using NCBI, by targeting 116 bp of the mitochondrial COI gene. Once PCR amplicons were identified for suitable detection, *in vitro* validation was performed using extracted DNA of *E. japonicus* and the non-target copepod *Mesocyclops* spp. from the water samples collected in Lake Biwa to evaluate primer-probe specificity. Using the *E. japonicus* primer-set, we did not detect PCR amplicons from the non-target species. The validation was also performed using DNA extracted from the 2017LB5 core samples at depth layers of 9.5 cm, 19.5 cm, and 29.5 cm. Among the qPCR amplicons of the 2017LB2 core samples, the 12 samples randomly collected were directly sequenced following ExoSAP-IT (USB Corporation, Cleveland, OH, USA) treatment. Sequences were determined using a commercial sequencing service (Eurofins Genomics, Tokyo, Japan).

### qPCR

DNA samples were quantified via real-time TaqMan quantitative PCR using the QuantStudio™ 1 Real-Time PCR System (Thermo Fisher Scientific, USA) in the Center for Marine Environmental Studies, Ehime University. The qPCR protocol followed that of Tsugeki *et al*. (2022). Briefly, the 1uL TaqMan reaction contained 900 nM of each forward and reverse primer, 125 nM TaqMan-Probe, and 10 µL qPCR master mix (TaqPath; Thermo Fisher Scientific) added to 4.0 µL template DNA solution. The final volume of the PCR was 20 µL after adding UltraPure™ DNase/RNase-Free Distilled Water (Thermo Fisher Scientific). The dilution gradient pattern of 20,000, 2,000, 200, and 20 copies per PCR reaction was proposed for the standard curve using the target DNA cloned into a plasmid. We ordered the cloning DNA of the 116-bp COI gene for qPCR amplification region using the plasmid to Eurofins Scientific (Tokyo, Japan).

The *R^2^* values of the regression analysis for the standard curves ranged from 0.983 to 0.993 (PCR efficiency = 81.2 to 98.4%). The qPCR data of the DNA copies (copies g^-1^ dry sed.) were shown as mean ± SD of four technical replicates in one sample. The limit of detection (LOD) of qPCR was six copies per PCR reaction for *E. japonicus*, and qPCR replicates that were not detected (less than the LOD) were treated as zero (0). One no-template control reaction (NTC) was included as the negative control in each run.

### Inhibitor test

To test the PCR inhibition effect of substances and minerals included in sediments, spike tests were performed for the 2017LB2 and 2017LB5 core samples (Fig. 1b). Detailed information on the spike test was provided in a previous study (Tsugeki *et al*., 2022) and ΔCt ≥ 3 cycle appeared to be evidence of inhibition (Hartman *et al*., 2005). We used the internal positive control (IPC; 207-bp, Nippon Gene Co. Ltd., Tokyo, Japan, an artificial DNA sequence designed to differ from any sequence of the organisms) as the spike DNA. Briefly, 1 µL plasmid, including the IPC (100 copies per PCR reaction), was added to the PCR template with the same PCR mixer as described above. We used the following primer and probe sets for IPC:

IPC1-5′: CCGAGCTTACAAGGCAGGTT
IPC1-3′: TGGCTCGTACACCAGCATACTAG
IPC1-Taq: [FAM] TAGCTTCAAGCATCTGGCTGTCGGC [TAMRA]

### Demographic traits and experimental test to explore potential sources of sedDNA

To determine the source of *E. japonicus* sedDNA, we evaluated the correlation between the sedDNA concentration and demographic traits of this copepod during recent decades, including biomass, specific growth rate, population production, and the number of subitaneous and resting eggs (Fig. 1g, h). Long-term monthly datasets of biomass, specific growth rate, and population production can be found in our previous reports (Liu *et al*., 2020b, 2021). The calculations of these population metrics are summarized in Table S4, with more details provided in previous studies (Liu *et al*., 2015, 2020b, 2021). Briefly, biomass (g dry weight m^-2^) was calculated as the product of the population density (individuals m^-2^) with the mean body dry weight (g individual^-1^). Specific growth rate (g day ^-1^) was estimated using a multiple regression equation with two parameters: mean water temperature (°C) during the copepod development and the food index as a proxy for availability of food required for *E. japonicus* growth (Liu *et al*., 2015, 2021). The food index was calculated from the ratio of the prosome length of the females collected from Lake Biwa to the expected prosome length of the females obtained from laboratory experiments under adequate food conditions. Population production (gC m^-2^ month^-1^) was calculated as the product of biomass and specific growth rate. The concentration of subitaneous eggs (eggs m^-2^) was estimated as the product of proportion of ovigerous female (%), ovigerous female density (individuals m^-2^), and clutch size (eggs clutch^-1^). Here, all parameters used in the calculation of subitaneous eggs were based on counts of individuals collected by field survey (Dur *et al*., 2022). The concentration of resting eggs (eggs g^-1^ dry sediment) was calculated as the ratio of the number of resting eggs counted by microscopy from each layer of the 2013LB core (eggs sample^-1^) to the dry weight for each layer of sediment (g dry sediment) (Liu *et al*., 2020a).

To determine whether *E. japonicus* sedDNA concentrations were regulated by DNA derived directly from resting eggs included in the analytical sediment, we divided the sediment sample into two fractions using a sieve to exclude the eggs (Fig. 1e). Since the minimum size of the resting egg of this copepod is approximately 80 µm (Liu *et al*., 2020a), the analytical sediments for DNA extraction were divided into two size classes using a 38-µm mesh-sieve on three randomly selected-layer samples (specifically, 2017LB5-2.5 cm, 2017LB5-10.5 cm, and 2017LB5-19.5 cm). Furthermore, to test the possibility of vertical movement of *E. japonicus* sedDNA through pore waters, we examined the sedDNA concentration in pore water and its residual sediment by separating them (Fig. 1f). The DNA extractions of the aforementioned samples were performed using previously reported protocols (Tsugeki *et al*., 2022).

### Possible factors regulating *E. japonicus* sedDNA

To explore potential factors regulating temporal variation in sedDNA concentrations, we analyzed chlorophyll-a pigments and algal remains. Sedimentary pigments of chlorophyll-a were investigated for the 2017LB2 core, and algal remains were investigated for the 2017LB7 core (Fig. 1d). Details of the method used for chlorophyll-a and algal remains are described in previous studies (Tsugeki *et al*., 2021, 2022).

## Data analysis

All statistical analyses were performed using R ver. 4.1.2 (R Core Team 2021) with the package “smatr” ver. 3.4.8. Type II regression models along with the standardized major axis method were used to determine the relationship between sedDNA concentrations of *E. japonicus* and demographic properties of biomass, specific growth rate, production, and abundance of *E. japonicus* subitaneous, resting eggs, and algal proxies. Since sedDNA and resting egg analysis for *E. japonicus* (Liu *et al*., 2020a) were conducted on different sediment cores (2017LB2, 2017LB5, and 2013LB; Fig. 1b, h), the chronological age of each analytical sample was slightly different. Therefore, prior to statistical analysis, each sedDNA and resting egg concentration in the respective core was converted to the value at 0.1-year intervals by linear interpolation using the AnalySeries 2.0.3 software package (Paillard *et al*., 1996). We then recalculated the average values within the year corresponding to the sedimentation period in each sample of the chronology 2017LB7 core.

The monthly data of demographic traits in *E. japonicus* including biomass, specific growth rate, production, and abundance of subitaneous eggs were annually averaged and converted to the value at 0.1-year intervals by linear interpolation. Thereafter, we calculated the average values in the respective sedimentation period of the 2017LB7 core samples, as mentioned above. These demographic data were limited to the year 2010. However, we estimated that the value until 2011 corresponded to the estimated age at 4.5 cm depth of 2017LB7 core by linear extrapolation using AnalySeries 2.0.3 software package (Paillard *et al*., 1996). Prior to Type II regression analysis, we tested the normality of the data with the Shapiro–Wilk test (significant level *p* < 0.05). The Shapiro–Wilk test rejected the hypothesis that sedDNA concentration and subitaneous eggs were normally distributed. Therefore, the data were log_10_-transformed to perform a Type II regression. We then used Type II regression models to determine the relationship between the log_10_-transformed values of *E. japonicus* sedDNA concentration and their demographic traits and algal proxies.

## Results

### sedDNA concentration of *E. japonicus*

The results of qPCR analysis in both cores (2017LB2 and 2017LB5; Fig. 1b) showed that *E. japonicus* sedDNA copies were detected continuously throughout the study period (Fig. 2a, b), except for three missing layers for the 2017LB5 core (5.5 cm, 6.5 cm, and 7.5 cm; Table S2). The mean DNA copy number of *E. japonicus* varied depending on the core depth and ranged from 6 ± 11 (mean ± SD) copies g^-1^ dry sediment (hereafter copies g^-1^) in the depth of the 2017LB2-28.5 cm layer (estimated age: 1924.5 ± 39.0) to 290,330 ± 30,848 copies g^-1^ in the 2017LB2-4.5 cm layer (estimated age: 2007.3 ± 0.4). Table S2). The sedDNA concentration of *E. japonicus* was high in the 3–4 replicates of samples in the upper 8.5 cm layers (estimated age: 1993.8 ± 1.0 and 1999.0 ± 0.7 for the 2017LB2 and 2017LB5 cores, respectively; Fig. 2a, Table S2). The DNA copy numbers below the 8.5 cm were relatively low but were detected in 1–4 replicates from all of the layers (Fig. 2b, Table S2).

**Figure 2.**
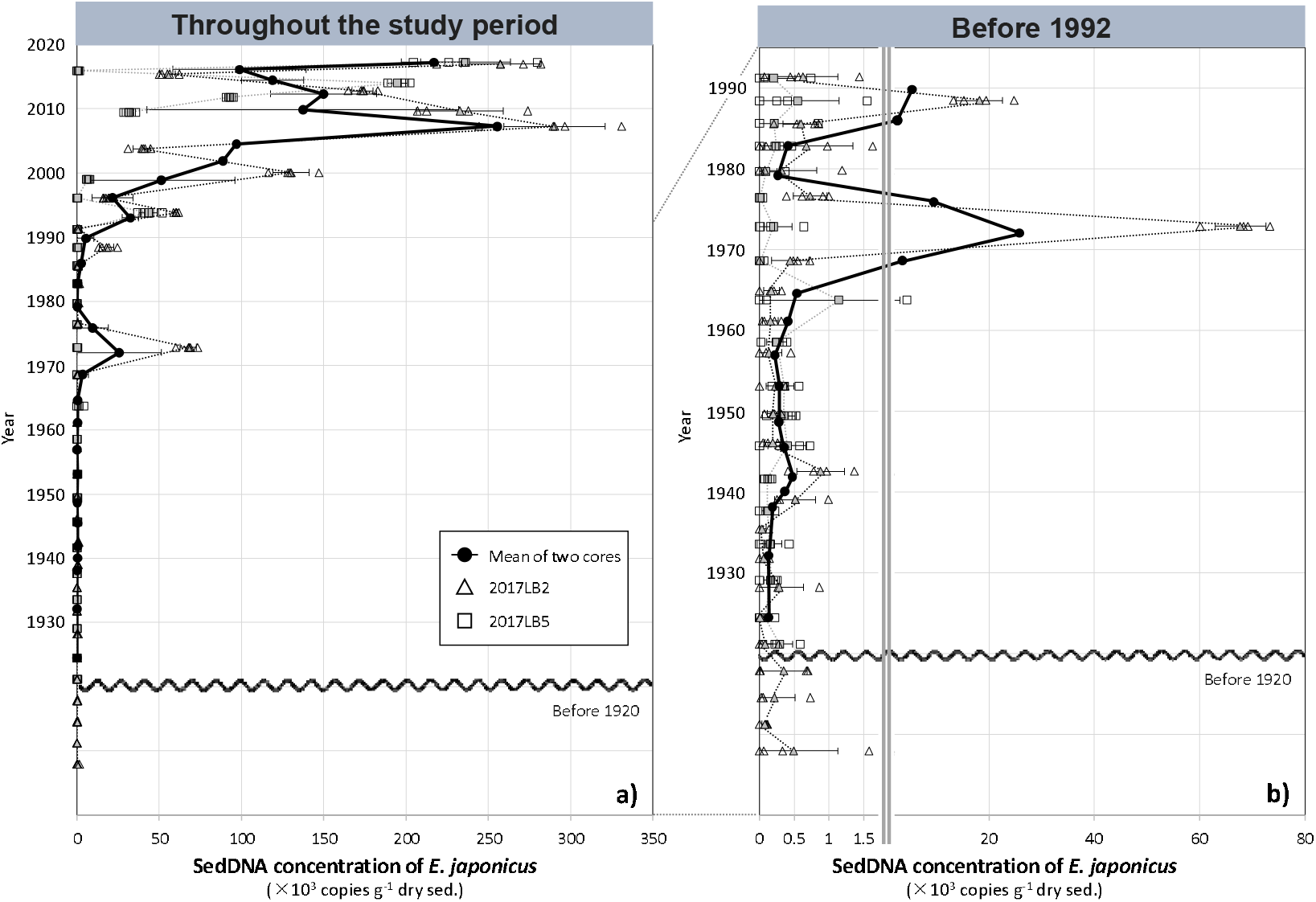
Temporal changes in *Eodiaptomus japonicus* sedimentary DNA concentration (gene copies of COI, g^-1^ of dry sediments): (a) observed throughout the study period, and (b) limited to the period before 1992. For both panels, the open triangles and squares indicate raw data of the 2017LB2 and 2017LB5 cores, respectively. The gray triangles and squares indicate the average mean in four replicates of 2017LB2 and 2017LB5 core samples, respectively. In panels (a) and (b), the black circles and thick solid lines with error bars indicate mean (*n* = 2) with SD of the 2017LB2 and 2017LB5 core samples within the year corresponding to the sedimentation period in the chronology 2017LB7 core.

No DNA of *E. japonicus* was detected in any of the negative controls for sub-sampling, DNA extraction, or PCR analysis, or the NTC, indicating that sample contamination did not occur during these processes. We also confirmed that only *E. japonicus* DNA was amplified through direct sequencing of PCR amplicons, using qPCR analysis of *E. japonicus* DNA.

A spike test was performed on both sediment core samples (2017LB2 and 2017LB5) for evaluating the effect of qPCR inhibition (Fig. 1b). All ΔCt values were below 1 (–1.0 to 0.87: 2017LB2, –0.49 to 0.21: 2017LB5; Table S3), indicating no inhibition.

### Long-term variation in demographic traits of *E. japonicus*

All the mean values of demographic traits such as biomass, specific growth rate, and production for *E. japonicus* were calculated depending on the time resolution of deposition period on each sample (see details in data analysis). The average biomass of *E. japonicus* till approximately the year 2000 was relatively low with a range of 0.7 to 1.2 g dry weight m^-2^, except for the values measured around 1970 (1.7 g dry weight m^-2^), but the average biomass rapidly increased approximately two-fold after 2010 compared to that before 2000 (Fig. S2a). Likewise, the average mean of their production was stable before approximately 2000, at less than 1.5 g C m^-2^, and then continuously increased up to 2010 (Fig. S2c). Contrarily, the average mean of specific growth rate (*g*) remained relatively constant after 1970 at approximately 0.07 day^-1^. The lowest value of specific growth rate was observed around 1970 and the highest value was observed around 2010 (Fig. S2b). The average mean of subitaneous eggs remained constant, being less than 1.5×10^5^ eggs m^-2^ up to approximately 2000, excluding the marked peak in the late 1990s, and continuously increased from 2000 onwards (Fig. S2d). The average mean of resting egg concentration recovered from sediment observation increased markedly from the mid-1980s, peaked around the early 1990s, and then decreased and stabilized (Fig. S2e).

### Correlations between *E. japonicus* sedDNA and its demographic traits with the concentrations of algal remains

A highly significant correlation was detected between the *E. japonicus* sedDNA concentration and *E. japonicus* production from 1971 to 2010 (Type II regression model: *R^2^*= 0.60, *p* = 0.001, *n* = 14; Fig. 3c), in addition to its biomass (*R^2^* = 0.51, *p* = 0.004, *n* = 14; Fig. 3a). A significant correlation was also detected between the sedDNA concentration and number of subitaneous *E. japonicus* eggs (*R^2^*= 0.52, *p* = 0.005, *n* = 13; Fig. 3d). In contrast, no significant relationship was observed between the sedDNA concentration and the specific growth rate (*R^2^* = 0.02, *p* = 0.66, *n* = 14; Fig. 3b), or the number of resting eggs (*R^2^* = 0.25, *p* = 0.066, *n* = 14; Fig. 3e). *Eodiaptomus japonicus* sedDNA concentrations were positively correlated with diatom valves (*R^2^* = 0.65, *p* < 0.001, *n* = 27; Fig. S4a), chlorophyll-a (*R^2^*= 0.71, *p* < 0.001, *n* = 29; Fig. S4b) and green algal cells (*R^2^* = 0.37, *p* = 0.001, *n* = 25; Fig. S4c), although these long-term trends differed from the trend of variation in *E. japonicus* sedDNA concentrations (Fig. S4d, e, f).

**Figure 3.**
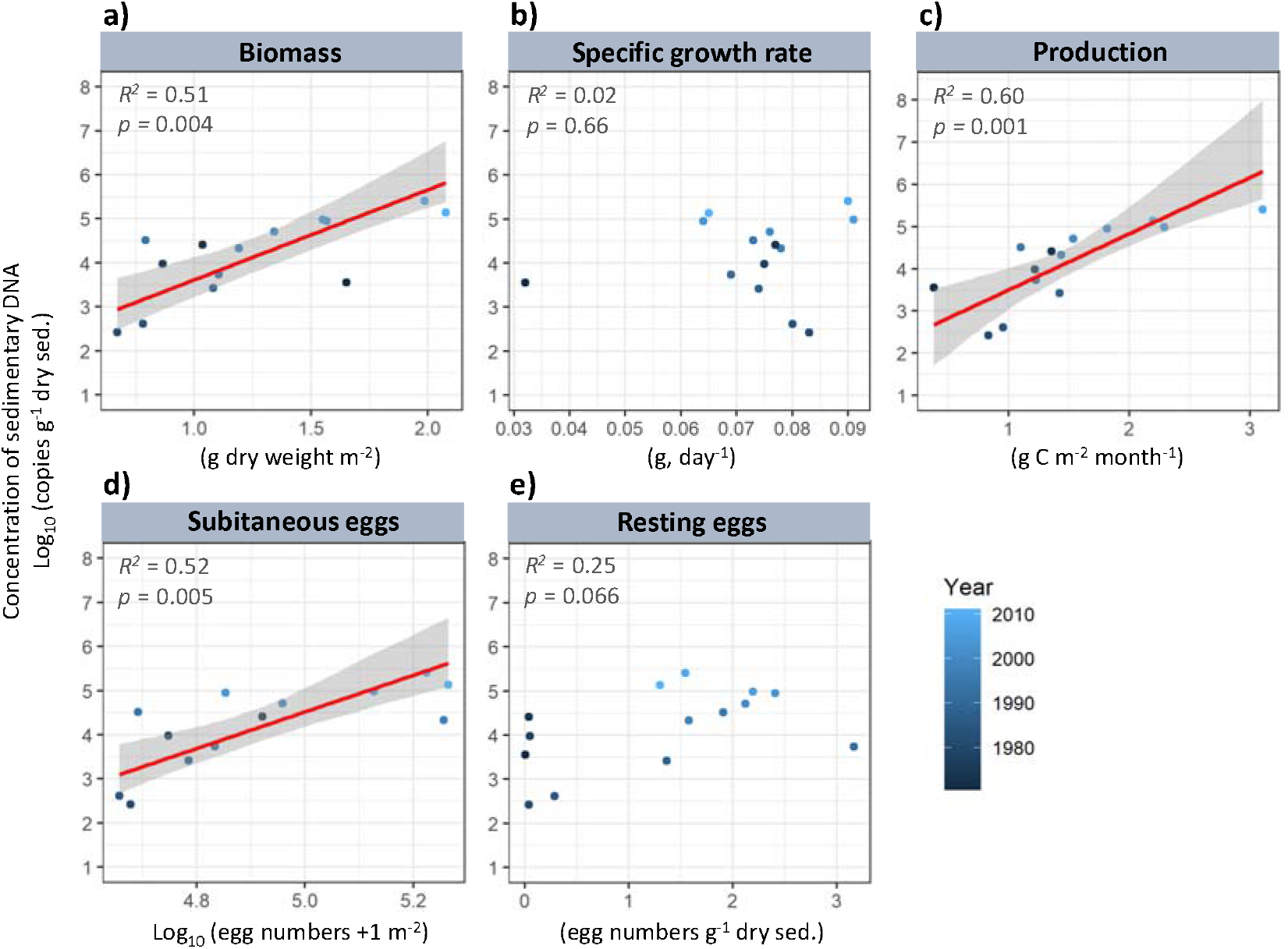
Relationships among log_10_-transformed sedimentary DNA concentration of *Eodiaptomus japonicus* (gene copies of COI, g^-1^ of dry sediments) and its biomass (g dry weight m^-2^; a), specific growth rate (day^-1^; b), production (g C m^-2^ month^-1^; c), log_10_-transformed subitaneous eggs (×10^3^ egg numbers m^-2^; d), and resting eggs (egg numbers g^-1^ dry sed.; e). The red line denotes a Type II regression model with a 95% confidence interval indicated by the gray zone.

### Potential sources of *E. japonicus* DNA

To explore the possibility of the vertical movement of sedDNA through pore waters, we examined the *E. japonicus* sedDNA concentration in pore water separating it from its residual sediment on three randomly collected samples. The qPCR did not detect *E. japonicus* environmental DNA (eDNA) from the pore water of all three samples from the 2017LB5 core (Fig. 1f, Table 1). In contrast, *E. japonicus* eDNA was detected in the residual sediment samples. Additionally, to assess the contribution of DNA derived directly from resting eggs included in the analytical sediment, we divided the sediment sample into two groups: one sieved to exclude the eggs and the other unsieved. *Eodiaptomus japonicus* sedDNA was detected in both sieved and unsieved groups, except for the sieved sample of 2017LB5-20 (Table 2). High concentrations of sedDNA were detected in the unsieved group for two of the three samples, namely 2017LB5-3 (11,630 ± 5,921 copies g^-1^) and 2017LB5-20 (1,449 ± 1,977 copies g^-1^), compared to those in the sieved samples. However, for 2017LB5-11, the concentration of sedDNA in the sieved sample (2,630 ± 4,555 copies g^-1^) was higher than in the unsieved sample (1,838 ± 3,184 copies g^-1^), although the difference between both values was minimal.

**Table 1.** The sedimentary DNA concentration of Eodiaptomus japonicus (gene copies of COI, g^-1^ of dry sediments) for pore water and its residual sediment samples.

| Sample ID (estimated age in sample depth and its age error; yr) | Core depth |  | Pore water sample | SD | Detection / total replicates | Residual sediment sample | SD | Detection / total replicates |
| --- | --- | --- | --- | --- | --- | --- | --- | --- |
|  | Top (cm) | Bottom (cm) | (mean copies g<sup>-1</sup> dry) |  |  | (mean copies g<sup>-1</sup> dry) |  |  |
| 2017LB5-3 (2014 ± 0.1) | 2 | 3 | ND | - | 0/4 | 725,030 | 98,992 | 4/4 |
| 2017LB5-11 (1994 ± 1.3) | 10 | 11 | ND | - | 0/4 | 2,665 | 608 | 4/4 |
| 2017LB5-20 (1964 ± 9.3) | 19 | 20 | ND | - | 0/4 | 12 | 21 | 1/4 |

**Table 2.** The sedimentary DNA concentration of Eodiaptomus japonicus (gene copies of COI, g^-1^ of dry sediments) in the sieved and unsieved sediment samples.

| Sample ID (estimated age in sample depth and its age error; yr) | Core depth |  | Sieved samples | SD | Detection / total replicates | Unsieved samples | SD | Detection / total replicates |
| --- | --- | --- | --- | --- | --- | --- | --- | --- |
|  | Top (cm) | Bottom (cm) | (mean copies g<sup>-1</sup> dry) |  |  | (mean copies g<sup>-1</sup> dry) |  |  |
| 2017LB5-3 (2014 ± 0.1) | 2 | 3 | 5,246 | 6,531 | 2/4 | 11,630 | 5,921 | 4/4 |
| 2017LB5-11 (1994 ± 1.3) | 10 | 11 | 2,630 | 4,555 | 1/4 | 1,838 | 3,184 | 1/4 |
| 2017LB5-20 (1964 ± 9.3) | 19 | 20 | ND | - | 0/4 | 1,449 | 1,977 | 2/4 |

## Discussion

Several studies have reported the long-term preservation of copepod eDNA in aquatic sediments (Bissett *et al*., 2005; Coolen *et al*., 2013; Xu *et al*., 2011). For example, Epp *et al*., (2015) detected copepod eDNA from sediment layers as old as around 10,000 to 7,500 years in a Greenland lake. However, continuous, and quantitative detection of copepod sedDNA has not yet been reported (Domaizon *et al*., 2017). To the best of our knowledge this is the first study to successfully detect and quantify sedDNA specific to an important copepod species, and to demonstrate the potential of such approaches for reconstructing past population dynamics with high temporal resolution.

The temporal variation in the concentration of *E. japonicus* sedDNA correlated significantly with its production and biomass, obtained via microscopic observations of regular field samples. In support of this, the number of reads for copepod DNA obtained by metabarcoding genetic analysis correlated with their biomass in water samples (Yang *et al*., 2017) although a meaningful relationship between eDNA abundance and biomass for some copepod species was not clearly observed (Djurhuus *et al*., 2018). Furthermore, a temporal increase in *E. japonicus* sedDNA concentration (more than 50-fold) was observed in the 1970s, when eutrophication occurred in Lake Biwa (Hyodo *et al*., 2008). Concordant with this finding, regular survey reports indicated the density of *E. japonicus* in Lake Biwa increased sharply in 1975, with densities more than 10-times higher than those in 1974 (Shiga Prefecture, 1964-1979). This coincidence of increased sedDNA concentration and the marked expansion of *E. japonicus* in the water column provides evidence that copepod sedDNA concentration can be a powerful tool to estimate their population biomass over several decades.

The temporal variation in the concentration of *E. japonicus* sedDNA was unlikely to be a result of degradation except for the top layer. Leaching from sediments has been observed mainly in coarse-textured sediments that enable fluid flow through layers (Andersen *et al*., 2012; Haile *et al*., 2007) and leaching is generally assumed to be a minor process in water-saturated sediments (Giguet-Covex *et al*., 2014), such as the fresh sediments investigated in the current study. Indeed, we did not detect *E. japonicus* eDNA in the pore water samples, as has previously been shown in studies on sedDNA of fish (Kuwae *et al*., 2020) and *Daphnia* (Tsugeki *et al*., 2022). Therefore, it is reasonable to conclude that leached *E. japonicus* eDNA only negligibly influences sedDNA. Although sedDNA might be degraded with time (Anderson-Carpenter *et al*., 2011), a recent experimental study showed almost no reduction in its concentration after an initial decline within the first month (Ogata *et al*., 2021). The concentrations of *E. japonicus* sedDNA in our study did not always show increasing trends toward the surface layers. Rather, it decreased toward the surface after the peak observed before 2010, except for the top surface layer. This indicates that temporal variation in the sedDNA of *E. japonicus* cannot be simply explained by time-dependent diagenetic degradation. The sedDNA concentration of *E. japonicus* correlated with some other proxies, such as pigment concentration and algal diatom remains. However, the long-term trends were different from those of *E. japonicus* sedDNA, indicating that *E. japonicus* sedDNA concentration is not regulated by phytoplankton sinking and/or their debris aggregations.

We detected that the sedDNA concentrations of *E. japonicus* correlated with the concentration of its subitaneous eggs. Moreover, we observed high concentrations of sedDNA from the unsieved samples, likely including resting eggs, in two of the three samples compared to those in the sieved samples that excluded eggs. However, we did not detect a significant correlation with the number of resting eggs, which is likely because in Lake Biwa, subitaneous eggs (rather than resting eggs) regulate the spawning activity of *E. japonicus* (Liu *et al*., 2020a). The contribution of resting eggs to total *E. japonicus* eggs in the water column was expected to be < 0.3% (Liu *et al*., 2020a). A recent study demonstrated that eDNA was released during spawning periods (Takeuchi *et al*., 2019; Tillotson *et al*., 2018). In addition, in a report of zooplankton *Daphnia* sedDNA, substances released during spawning activities were assumed to be a significant source of DNA archived in sediments (Tsugeki *et al*., 2022). Therefore, these findings emphasize the importance of sedDNA-sourced material being released during the main spawning period. As shown in the present study, the sedDNA concentrations of *E. japonicus* increased more than 3-fold during the 2000s compared to those before the 1990s, which was significantly correlated with their production and biomass. The production of *E. japonicus* increased more than double in the 2000s, mainly due to the decline in predation effects (Liu *et al*., 2020b, 2021), which occurred in parallel with an increase in their specific growth rate and number of subitaneous eggs. These facts indicate that copepod sedDNA can be applied as a powerful tracking tool for the reconstruction of their production and biomass related to the intensity of spawning activity.

In conclusion, we quantitatively and successfully detected *E. japonicus* sedDNA over a 100-year period, demonstrating the potential utility of sedDNA in evaluating the long-term quantitative dynamics of copepods. The absence of *E. japonicus* sedDNA in the pore water samples suggested that temporal variation in its sedDNA concentration was not interrupted by leaching reactions after sedimentation. The historical variation in sedDNA concentration was similar to its production, biomass, and subitaneous eggs found in regular *in situ* survey samples. Therefore, the *E. japonicus* sedDNA concentration is a possible indicator of the magnitude of productive activities in the water. Overall, we confirmed that sedDNA can be used as a proxy for the production and spawning activity of copepods over at least 100 years of sedimentary records.

## Acknowledgements

We appreciate two anonymous reviewers for their constructive comments and suggestions. We are also grateful to M. Honjo, M. Sakata, S. Goda, T. Akatsuka, and J. Kurata for their assistance with laboratory analysis and field sampling. We also thank Shiga Prefectural Fisheries Experiment Station for providing preserved zooplankton sample collection. This study was supported by Grants-in-Aid for Scientific Research (17K20045, 21K12273, 21H01170, and 18H03961) from the Japan Society for the Promotion of Science (JSPS) and partly supported by a special research grant from Matsuyama University; the Academic Research Organization Joint Usage/Research Grants from Leading Academia in Marine and Environment Pollution Research (LaMer), Ehime University; the Center for Ecological Research (2017–2021 jurc-cer), Kyoto University; the cooperative research program (17A065) of the Center for Advanced Marine Core Research, Kochi University, and the Environment Research and Technology Development Fund (JPMEERF20204004) of the Environmental Restoration and Conservation Agency, Japan; and grants from the Ministry of Agriculture, Forestry and Fisheries, Japan for a research project titled *Development of technologies for mitigation and adaptation to climate change in Agriculture, Forestry and Fisheries*.

## Data availability statement

The data that support the findings of this study are available from the corresponding author upon reasonable request.

## Author contributions

Conceptualization: N.T. and K.N. Fieldwork: N.T., M.K. Laboratory work: K.N., N.T., M.K. Design and testing of qPCR primers and probes: K.N., H.D. Preparation of figures and tables: K.N., N.T. Writing and data analysis: N.T., K.N., H.D., M.K., X.L., G.D., S.B.

## Conflict of interest

The authors declare no conflict of interest.

## Supporting information

Additional supporting information can be found online in the Supporting Information section at the end of this article.

## Materials & Correspondence

Correspondence to Narumi Tsugeki.

## Notes

### Competing Interest Statement

The authors have declared no competing interest.

### Summary of Updates

The section on the methods was updated to clarify; Figure 2 was revised; author affiliation was updated

